# Deep learning-driven morphology analysis enables label-free classification of therapeutic agent-naive versus resistant cancer cells

**DOI:** 10.1101/2025.01.22.634357

**Authors:** Cyril Ramathal, Kiran Saini, Zhouyang Lian, Christian Corona, Tiffine Pham, Ryan Carelli, Stephane C. Boutet, Manisha Ray, Sunantha Sethuraman, Vivian Prindle, Evelyn Lattmann, Maria-Grazia Molvetti, Andreas Dzung, Mitchell P. Levesque, Matt Barnes, Andreja Jovic

## Abstract

Therapeutic drug treatments of solid tumors are often undermined by various resistance mechanisms. Identification of drug-resistance phenotypes at the single cell level is challenging because conventional molecular methods are cell-destructive, labor-intensive, and cost-prohibitive. To overcome these challenges, we developed an orthogonal approach to drug-resistance phenotyping, through the use of deep-learning-driven morphology analysis of single, high resolution cell images.

Specifically, we trained deep learning-based drug resistance classifiers using cell images from 5 different cell lines that were rendered resistant to 5 different therapeutic agents, using a foundation model framework. With high accuracy, the classifier correctly predicted naive or resistance phenotypes across all cancer types and across all the therapeutic agent types (chemotherapeutic, targeted) tested. These results showed that morphology can capture complex phenotype information in the context of drug treatment. To demonstrate the potential clinical utility of the drug resistance classifier, it was applied to a dissociated tumor biopsy and the resulting phenotype predictions were in close concordance with scRNASeq analysis of the biopsy.

Our study highlights the potential of deep-learning-driven morphology analysis to provide complex phenotype information, and ultimately shape oncology drug treatment strategies at the patient-level in a clinical context.

## Introduction

Targeted therapeutic and chemotherapeutic agents have been developed to ablate the tumor cell population within solid tumors. Recent examples of targeted therapeutic agents include vemurafenib^1^ and binimetinib^2^, which target the BRAF- and MEK-driven signaling pathways contributing to the molecular etiology of melanoma, among other tumor types. Meanwhile, chemotherapeutic agents including cisplatin, doxorubicin, and paclitaxel, have been used in a non-cell selective manner to ablate the tumor cell population in patients ailing from various types of cancers, including lung and ovarian cancers. While these therapeutic agents have exhibited durable responses in subsets of patients, various resistance mechanisms have been uncovered that undermine treatment. These resistance mechanisms can be broadly categorized as intrinsic and acquired^3–5^ where the former case refers to underlying genetic or non-genetic mechanisms that exist before a treatment has been applied. The latter refers to molecular or biochemical changes that may develop in the tumor cell as an ‘escape’ mechanism from the selective pressure of the therapy over time.

Various methods have been applied to uncover resistance mechanisms to therapies and to ultimately define resistant phenotypes at the whole population and single cell level. Since phenotyping provides a readout of a tumor’s response or potential response to therapies, it can be used to direct the course of treatment options in a patient-centric manner^6^. For example, gene expression and proteomics have been used to identify resistance phenotypes in various cancer types, enabling a deeper understanding of therapeutic response and providing insight into additional treatment strategies^7^. However, many phenotyping methods are end-point in nature precluding further sample analysis, and are time-consuming, labor-intensive, cost-prohibitive, and require cell labeling. Therefore, approaches are needed that address these limitations.

Morphology analysis has recently emerged as a powerful means of assessing cellular phenotypes in the context of therapeutic drug discovery and drug response^8^. We hypothesized that morphology analysis could enable label-free classification of drug response phenotypes. To this end, we used the Deepcell platform^9,10^, which combines microfluidics, high resolution brightfield imaging, and deep-learning within a foundation model framework to provide quantitative single cell image analysis^11^.

The goal of our study was to use morphology analysis to develop a drug resistance phenotype classifier to enable label-free phenotyping of cancer cells at the single-cell level. The classifier was used to distinguish between drug naive versus drug resistant cells across various cancer types and therapeutic agents (i.e. chemotherapeutic and targeted agents) (Fig. 1). In addition, as a proof-of-concept to demonstrate clinical applicability, we employed the classifier to phenotype a dissociated biopsy sample and used single cell RNA Sequencing (scRNASeq) to assess the percentage of drug-resistant cells for comparison. This study demonstrates that morphology can accurately capture complex phenotype information in the context of drug response. We envision that when combined with sorting capabilities, our morphology analysis will enable isolation of viable, unlabeled cells for further downstream or functional analyses.

**Fig. 1:**
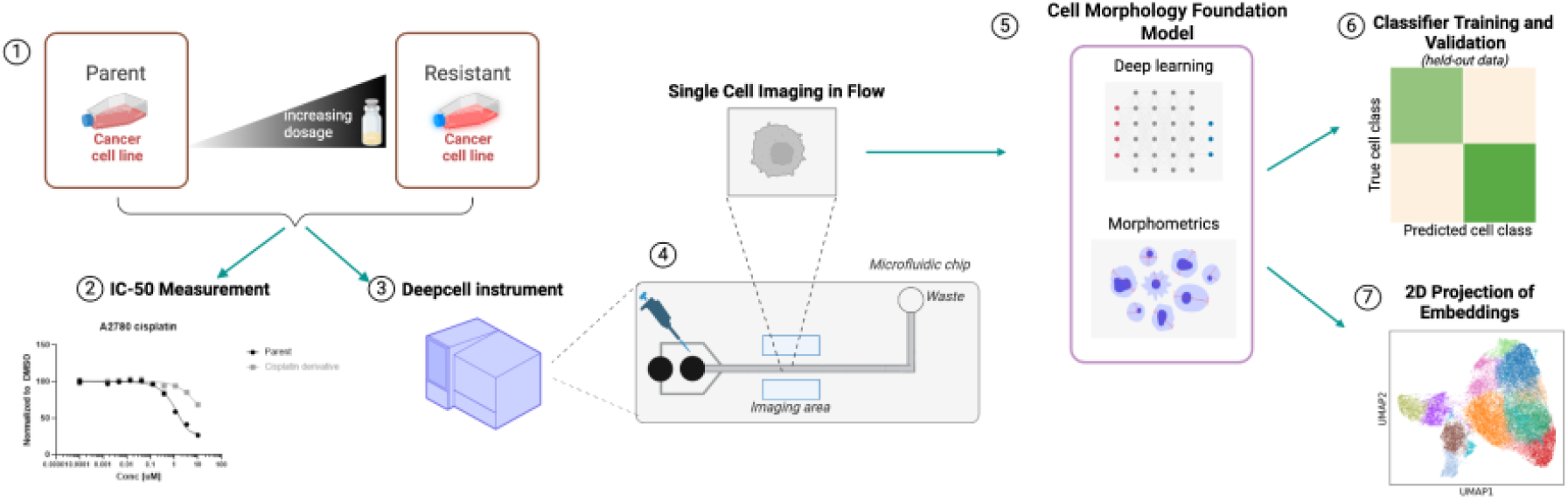
Schematic of the workflow for training and applying the Cell Morphology Foundation Model-based classifier for predicting drug resistance. 1) Creation of drug resistant cell lines, 2) verification of resistance phenotype through IC50 measurements, 3) processing and 4) imaging of naive and resistant cells on the Deepcell instrument, 5) extraction of morphology features using the Cell Morphology Foundation Model, 6) Classifier training and validation based on the images of pure populations of naive and resistant cell lines, and 7) projection of extracted morphological features in 2D for visualization and analysis of morphology patterns.

More generally, for the field of clinical oncology, this approach has the potential to provide patient-level assessments of therapeutic efficacy in a cost-effective and label-free manner.

## Materials and Methods

| Sample Name | Vendor/Source | Cancer Type | Catalog No. |
| --- | --- | --- | --- |
| A2780 | ECACC | Ovarian Cancer | 93112519 |
| A2780<br>Cisplatin-Resistant |  |  | 93112517 |
| A2780<br>Doxorubicin-Resistant |  |  | 93112520 |
| NCI-H460<br>(H460) | ATCC | Lung Cancer | HTB-177 |
| M010817 | Univ. of Zurich | Melanoma | N/A |
| M130425 | Univ. of Zurich | Melanoma | N/A |
| WM983B | Wistar Institute | Melanoma | N/A |
| DTC | Discovery Life Sciences | Lung Cancer | BTC1000-H1120571225091416MS |

Generation of chemotherapeutic resistant cells (NCI-H460, A2780)

### Generation of NCI-H460 Paclitaxel resistant cells

To generate NCI-H460 (H460) paclitaxel-resistant cells, cells were cultured with increasing concentrations of paclitaxel over the span of 3 months. H460 were split and allowed to attach to flask overnight before a dose of paclitaxel was added to the media and allowed to grow in the presence of paclitaxel for a few days. The cells were initially cultured at a low dose (0.5 nM paclitaxel). Over subsequent passaging of the cells, the dose of paclitaxel was gradually increased until reaching a concentration of 200nM (∼10x the IC50 of the parental line).

*Generation of targeted-therapy resistant melanoma cells (M010817, M130425, WM983B)* Melanoma cells were seeded in T25 flasks and cultured in RPMI-1640 medium (Gibco, cat. #R8758) supplemented with 5 mmol/L glutamine (Gibco, cat. #25030-081), 1 mmol/L sodium pyruvate (Sigma-Aldrich, cat. #S8636), 1% penicillin-streptomycin (Gibco, cat. #15070-063), and 10% heat-inactivated fetal bovine serum (FBS, Lonza) and maintained at 37°C and 5% CO2. Once 60-80% confluence was reached, cells were treated twice a week with binimetinib (Selleckchem, Cat.No. S7007), starting with a drug concentration of 10 nM. Drug concentrations were increased by approximately 50 nM each time the cells reached confluence and remained viable after passaging. The highest-used concentrations ranged between 500 to 1000 nM, depending on the tolerance of the cells.

### IC50 assays

A2780 and H460 cells were plated at 2,500 cells/well in 384-well plates (Corning, 3765) and dosed with compound (HP D300 Digital Dispenser). Cell viability was assessed on day 3 using CellTiter-Glo 2.0 Cell Viability Assay according to manufacturer’s protocol (Promega, G9241) and luminescence was quantified using a PerkinElmer Envision (2104 Multilabel Reader).

Melanoma cells were seeded into 96-well plates and treated 1 day after seeding for 3 days with increasing concentrations of the indicated drug, ranging from 1 nM to 10000 nM final concentration. Each condition was performed in triplicate. After 3 days of treatment, the medium was exchanged with a cell culture medium containing resazurin sodium salt (0.015 mg/ml, catalog number R7017, Sigma-Aldrich) and incubated for 2.5 hours at 37 °C. The fluorescence of metabolized resorufin that correlates with the number of living cells was measured using an excitation wavelength of 535 nm and an emission wavelength of 595 nm.

### Sample Preparation for Deepcell Instrument processing

Cryopreserved samples (cell lines, lung cancer DTC) were thawed in a water bath at 37°C until a small amount of ice was visible in the vial. Cells were then transferred to a 50 mL conical Falcon tube (Corning REF 430829), and 1mL of RPMI+ GlutaMAX (Gibco REF 61870-036) + 10% FBS (Peak Serum CAT 04A1232) was slowly added over the course of 1 minute to neutralize the freezing medium. An additional 8mL of medium at room temperature was added over the next minute, for a total volume of 10mL. The cell mixture was spun down at 300g for 5 minutes, and the cell pellet was resuspended in the Deepcell sample buffer to reach a concentration of 1e6 cells/mL. The resulting samples were imaged at ∼30 cells/s on at least two Deepcell instruments, and a minimum of 10,000 images were collected for each sample.

### Morphology-based Classifier Development and Implementation

The images from each sample (H460, A2780, M010817, WM983B, 130425) were annotated as “naive” or “drug-resistant” based on the sample phenotype. The DTC sample was annotated as “unknown”. Each image was encoded as a combination of morphometric features and deep learning-based features using a cell morphology foundation model. Low quality images such as out of focus cells and cell debris were filtered out from the data set. After filtering, equal numbers of images were sampled from the parent vs resistant populations for each sample.

These images were split into training and test sets based on the instrument used to image each sample replicate. A random forest classification model was trained on the image encoding to predict parent vs resistant type for each sample. The percentage of images from the test set classified as parent vs resistant was generated to calculate model metrics such as class confusion matrices. After filtering for out of focus and debris images, the percentage of images from the DTC samples classified as parent vs resistant was generated to compare against scRNASeq results.

### Foundation Model Development

The cell morphology foundation model is a hybrid architecture combining self-supervised learning embeddings and computer vision morphometrics. We implemented a variant of the VICReg method^12^ to construct the self-supervised deep learning embedding model. Computer vision algorithms were used to quantitatively extract morphometric features, such as cell shape, size, intensity, and texture. At the time of this study, the HFM training dataset included polystyrene beads, human cancer cell lines, immune cells, stromal cells, healthy and diseased tissue samples, and iPSC-derived differentiated cells.

### scRNASeq Processing and Analysis

Cells from the Lung Cancer DTC were prepared for scRNASeq processing by counting and resuspending them at a concentration of 1000 cells/uL in PBS supplemented with 0.04% BSA. Cells were run in duplicate using the 10x Chromium Next GEM Single Cell 3’ Reagent Kits v3.1 Dual Index (PN-1000268, 10x Genomics, Pleasanton, CA, United States) per manufacturer guidelines. 5000 cells were targeted for each library. The cells were then loaded into individual channels on the Chromium Next Gem Chip G (PN-1000127) and encapsulated into gel beads in emulsion (GEMs) via 10x Chromium iX instrument (PN-1000328). Single cell GEMs were lysed and underwent reverse transcription, resulting in barcoded cDNA. The resulting barcoded libraries underwent amplification, fragmentation, end-repair, adaptor ligation, and sample index attachment using Dual Index Kit TT Set A (PN-1000215). Final single cell transcriptome libraries were sequenced at a minimum depth of 20,000 reads per cell on the Illumina Nextseq 2000 and Novaseq 600 instruments as follows: 28 cycles (Read 1), 10 cycles (i7 Index), 10 cycles (i5 index), 90 cycles (Read 2).

The sequenced reads were processed using the Cell Ranger analysis pipeline v.7.0.1 (3) to generate FASTQ files and align sequencing reads to a pre-built human reference transcriptome version GRCh38-3.0.0 by 10x Genomics. The raw binary format output with cell barcodes and unique molecular identifier count results were used for the downstream analysis. Using automatic thresholding via median absolute deviations, the number of counts per barcode (count depth), the number of genes per barcode, and the fraction of counts from mitochondrial genes per barcode, low quality cells were removed. Ambient RNA and doublets were removed using soupX and scrublet correspondingly. The remaining raw counts were normalized via shifted logarithm. Batch correction was performed using highly variable genes. The subsequent high dimensional processed data was visualized with UMAP in 2-D space. Expression levels of the processed signature gene set (for paclitaxel resistance) were used to identify naive and paclitaxel-resistant cells.

The distribution of the sum of the expression values of the paclitaxel signature^13^ was used to identify threshold values with which to differentiate therapeutic-naive and resistant cells, respectively. The value 0 was chosen as the threshold value for classifying cells as naive or resistant, i.e. cells with the sum of paclitaxel-resistant expression values greater than 0 were classified as resistant and cells with values less than 0 were classified as naive. The sum of the expression levels was compared against the threshold values to determine the percentage of naive and resistant cells accordingly.

## Results

### Morphology analysis of therapeutic agent-resistant cell lines

We developed a morphology-based classifier to distinguish between therapeutic agent resistant and naive cells, using the Deepcell platform. We generated 6 resistant cell lines across three different cancer types using chemotherapeutic and targeted therapeutic agents (Table 1) and performed morphology analyses on each pair of resistant and naive cells. The workflow entailed processing single cell suspensions of the naive and resistant cells individually on the platform, extracting morphology features from each cell image, and training a classifier to distinguish between the two populations based on the extracted features (Fig. 1). IC50 curves were generated for each pair of resistant and naive cell lines, in order to verify the resistance phenotype (Suppl. Fig. 1).

**Table 1:** Table of the combinations of cell lines, cancer types, and therapeutic agents, and therapeutic agent types used for this study

| Cell Line | Cancer Type | Therapeutic agent | Therapy type |
| --- | --- | --- | --- |
| A2780 | Ovarian<br>Adenocarcinoma | Cisplatin | Chemotherapy |
|  |  | Doxorubicin |  |
| NCI-H460 | Lung Carcinoma | Paclitaxel |  |
| M010817 | Melanoma | Binimetinib | Targeted Therapy |
| 130425 |  |  |  |
| WM983B |  |  |  |

For each cell line pair, we developed a classifier to distinguish between naive and resistant cells, based only on morphology. For all 6 cell line pairs, we noted that the classifiers were able to distinguish the resistant cells from the parent cells with high accuracy, >=80% (Fig. 2, Suppl. Fig. 2). To gain further insight into the morphology differences between naive and resistant cells, the extracted morphometric and deep learning features for each cell were combined into a single vector (“embedding”) and plotted in 2-dimensions using Uniform Manifold Approximation and Projection (UMAP) (Fig. 2B); these plots portrayed that the majority of parent and resistant cells occupied unique parts of “morphology space”, reflected in the high accuracy classifier performance for each pair. Representative images of each cell population further highlighted these morphology differences (Fig. 2C). We then used Jensen-Shannon Divergence (“Divergence Score”) to uncover the top morphology features that distinguished naive from resistant cells for each pair (Fig. 2D, Suppl. Figs. 2C, G, K, O, S). This analysis revealed that a mixture of morphometric and deep learning features contributed to the differences between the two populations. Plotting of individual top features highlighted the distribution differences between parent and resistant cells (Fig. 2E, Suppl. Figs. 2D, H, L, P, T).

**Fig. 2:**
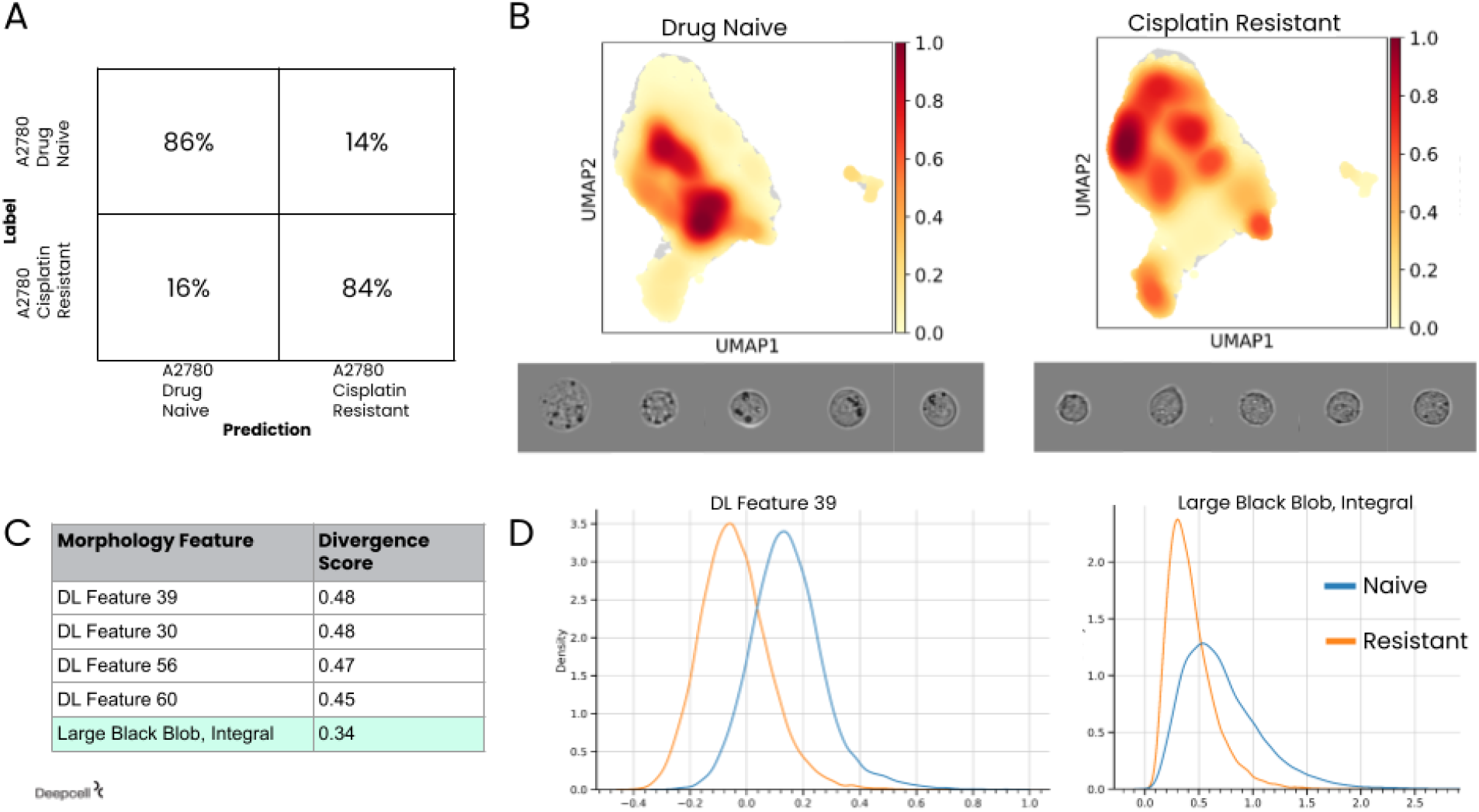
Morphology analysis of A2780 cisplatin resistant and naive cells reveals key features differentiating the two populations enabling high accuracy classification based on morphology alone. A) Confusion matrix of the classifier for predicting Cisplatin-resistant and naive A2780 cells based on morphology. B) UMAPs of the morphology features extracted by the Cell Morphology Foundation Model in density plot form, highlighting the morphological differences between the two populations. Representative images of each population beneath the respective UMAPs. C) Top 4 morphology features distinguishing the Cisplatin and Naive populations in A2780 cells, and the top morphometric feature in teal. D) Distributions of the top Deep Learning and Morphometric features between the naive and resistant populations.

For instance, the morphology features that distinguished A2780 (Ovarian Cancer) Cisplatin resistant from naive cells were a number of texture-based features, including “small black blobs” (Fig. 2C). Interestingly, texture-based features also distinguished Doxorubicin resistant A2780 cells from naive cells and Paclitaxel resistant H460 cells from naïve cells; for the former two instances, our analysis revealed that the resistant cells had fewer black blobs on their surfaces relative to naive cells (Suppl. Fig. 2C, G). In the case of the melanoma cell line M010817, the Binimetinib resistant cells also could be differentiated from the naive cells based largely upon pixel intensity features (Suppl. Fig. 2K). For melanoma cell line 130425, the size-based feature minimum feret was the morphometric feature that best distinguished the Binimetinib resistant vs. Naive cells, which confirmed the trend that was observed in the images of the two populations (Suppl. Figs. 2N, O). A similar increase in size was also observed in the Binimetinib resistant cells for the melanoma cell line WM983B (Suppl. Figs. 2R, S). For WM983B resistant cells, we also observed a subpopulation of pigmented cells (Suppl. Fig. 2R-red box), highlighting the morphological diversity in the resistant cell population.

In summary, our analysis demonstrated that morphology could be used to distinguish between naive and resistant cells with high accuracy across various cancer and drug types, providing insight into the key features distinguishing populations.

### Comparison of Morphology data across cell line pairs

In order to analyze how different perturbations affected the same cell line, we combined the morphology data sets from the ovarian cancer cell line A2780 into a heatmap, which entailed therapeutic naive cells and the Doxorubicin-resistant and the Cisplatin-resistant populations (Fig. 3A). The resulting heatmap provided an effective visualization to identify groups of features that exhibited similar or differing patterns across the three populations. As was noted in the previous section, the heatmap highlighted that a combination of texture (in green) and deep learning features (in blue) discriminated the resistant cells from the naive cells. Interestingly, the heatmap revealed that the two resistant populations were more similar morphologically to one another relative to the therapeutic-naive cells. These results show that while the respective cell lines are resistant to two different chemotherapeutics, there are instances of morphological overlap, which might indicate that similar molecular pathways might be engaged in the resistant cells.

**Fig. 3:**
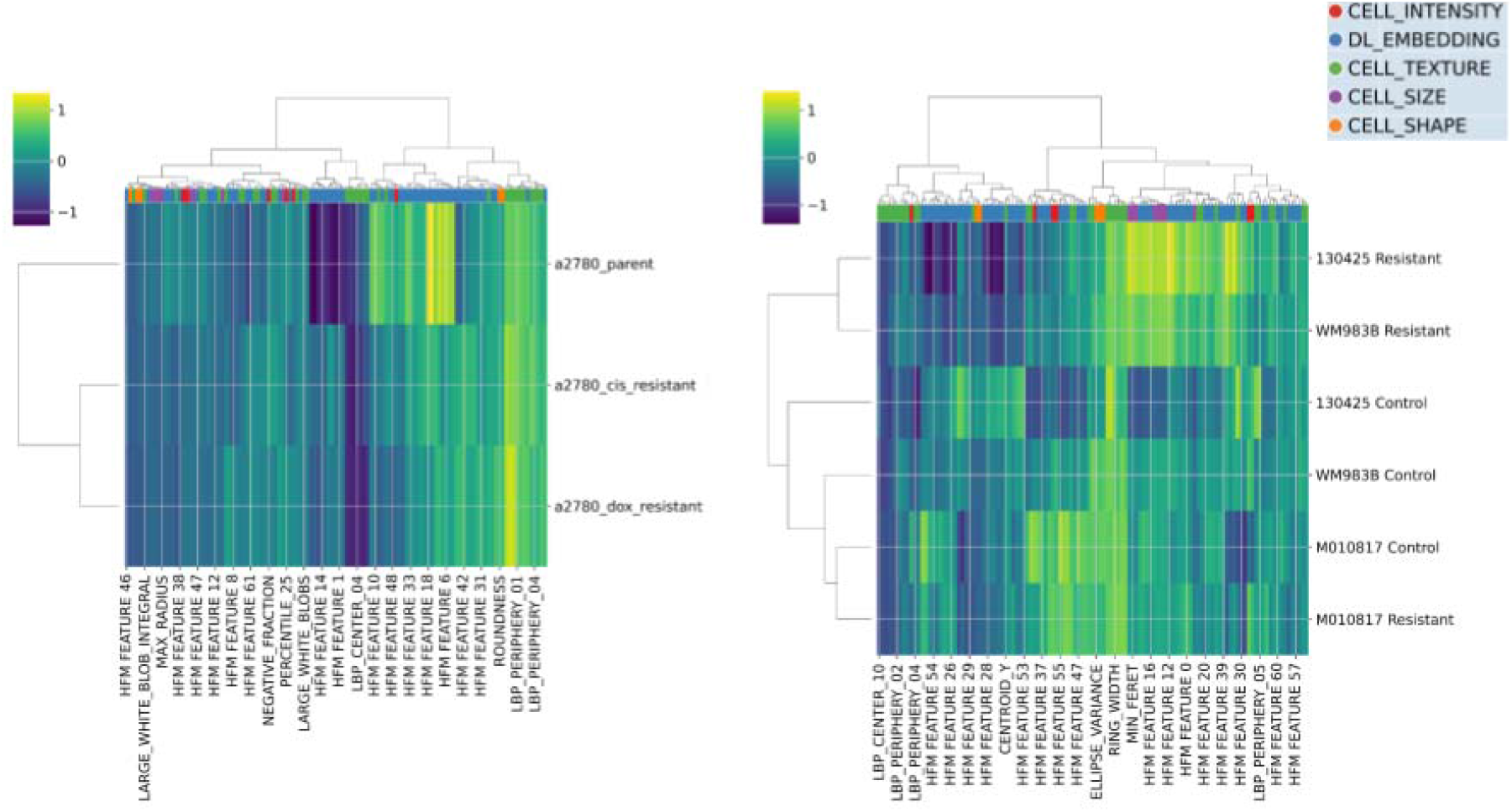
Heatmaps (with dendrograms) of the average morphology feature values for the same cell rendered resistant to different drugs (left), and a different set of cells rendered resistant to the same drug (right). (Left) A2780 cells rendered resistant to cisplatin are morphologically more similar to doxorubicin resistant cells relative to the naive cells. (Right) Within the melanoma cell lines, M010817 Binimetinib Resistant and naive are most similar to one another, as are 130425 Resistant and WM983B Resistant.

We also combined the morphology data from the 3 melanoma cell lines rendered resistant to the MEK inhibitor Binimetinib (M130425, M010817, and WM983B) in a heatmap (Fig. 3B), providing insight into how the same perturbation affects different melanoma cell lines. The heatmap showed that the M010817 naive and resistant cells were morphologically most similar to each other relative to the other populations. Heatmap analysis also revealed that M130425 naive cells and WM983B naive cells were most similar to each other relative to the rest of the cells; and similarly, M130425 resistant cells and WM983B resistant cells were most similar to each other relative to the rest of the cells in this group. Based on their morphological profiles, our results indicate that the cell lines WM983B and M130425 potentially overlap in terms of their mechanisms of Binimetinib resistance development; however, we did not observe a pigmented subpopulation in resistant M130425, which was present in resistant WM983B.

The ability to visualize morphology profiles of various cell lines and various perturbation conditions in heatmap form revealed various morphological patterns and trends, similar to standard approaches used for gene expression analysis of naive and resistant populations.

### Application of morphology classifier data to a clinical sample

As a proof-of-concept to demonstrate the applicability of our morphology classifiers to clinically-relevant samples, the H460 Paclitaxel-resistant classifier (Suppl. Fig. 2F) was applied to a lung cancer dissociated tumor cell (DTC) sample that was imaged on the Deepcell platform. In parallel, we collected scRNASeq data for this DTC to have a ground-truth estimate of the percentage of paclitaxel resistant cells in the sample based on an extant transcriptional signature^13^. The DTC was processed on the Deepcell platform and the H460 Paclitaxel Classifier was applied to the resulting morphology data to generate predictions of the percentage of Paclitaxel resistant and naive cells in the sample. The classifier predicted that approximately 35% of cells were Paclitaxel resistant in this sample, while scRNASeq analysis revealed that approximately 45% of cells were resistant based on the gene signature (Fig. 4A).

**Fig. 4:**
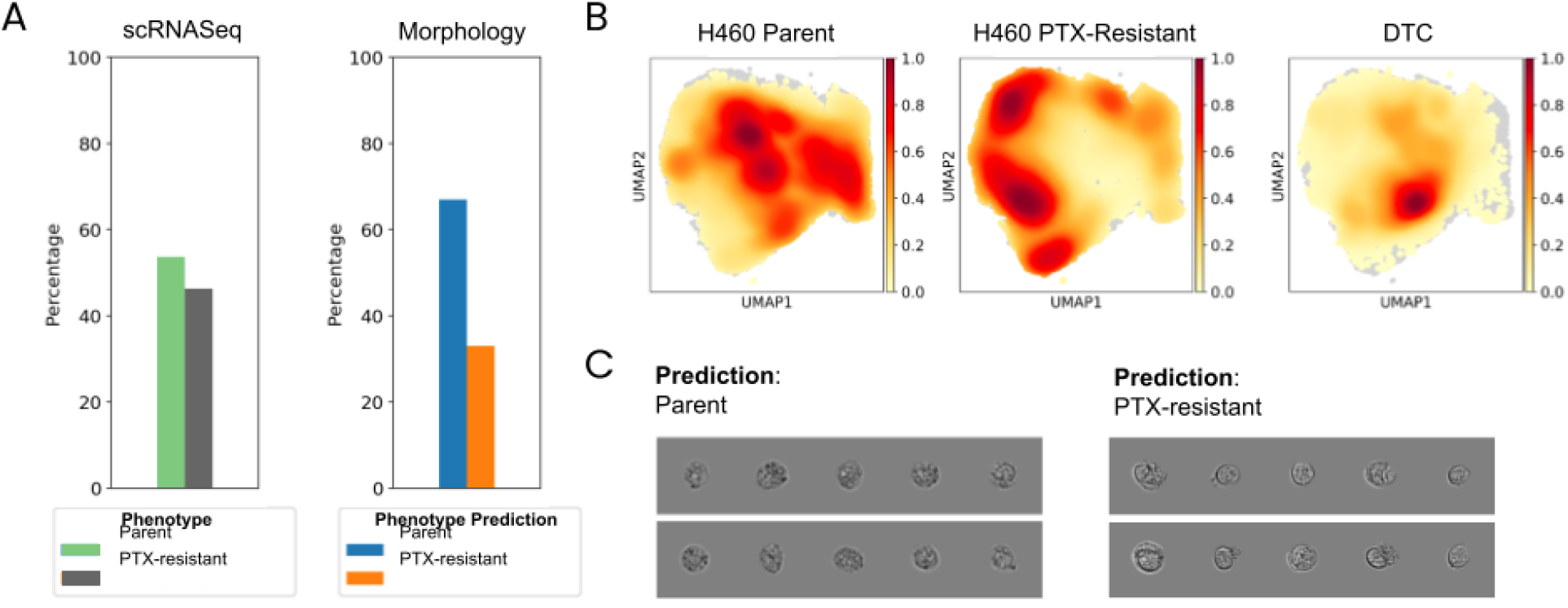
Applying the H460 Paclitaxel resistance classifier to a dissociated tumor cell (DTC) sample of a lung cancer patient for predicting the resistance phenotype, and comparing to phenotype data from transcriptional analysis. A) Transcriptional analysis of scRNASeq data from the DTC revealed that about half of the tumor cells are paclitaxel resistant based on a published gene signature (left) (). The morphology-based H460 Paclitaxel resistance classifier predicted about one third of cells are paclitaxel resistant in the same DTC (right). B) UMAPs of the extracted features from the H460 naive, H460 Paclitaxel-resistant, and DTC populations, which highlight how the DTC sample morphology data overlaps with each H460 population in morphology space. C) Representative images of predicted naive and paclitaxel resistant cells from the DTC sample.

We next used the embeddings generated from the DTC to create a UMAP of its morphological profile, in comparison to the H460 parent and resistant populations (Fig. 4B); this analysis corroborated the results highlighted by the classifier that the DTC was a mixture of naive and resistant cells. In addition, we noted morphology differences appreciable by eye according to representative images of predicted naive and resistant cells (Fig. 4C), which aligned with the morphology differences noted with the Paclitaxel naive and resistant H460 cells (Suppl. Fig. 2F). These results suggest that morphology-based classifiers provide valuable phenotype information for clinical samples, which can be used for predicting therapeutic response.

## Discussion

Acquired and innate resistance mechanisms to chemotherapeutic and targeted cancer therapies have presented a hindrance to the clinical response rates across cancer types. Various methods have been applied to uncover mechanisms of resistance to therapies and to ultimately define resistant phenotypes at the whole population and single cell level. While many of these methodologies have relied on molecular phenotyping of the drug-resistant population, a method to identify the resistant population of cells from the total population by label-free methods has remained elusive. While many of these methodologies have relied on molecular phenotyping of the drug-resistant population, a method to identify the resistant population of cells from the total population by cost-effective and non-labor-intensive methods has remained elusive.

Morphology analysis, also called image-based profiling, is a promising and rapidly developing approach for biological discovery. It has been applied to the various parts of drug discovery^8^, from screening^14^ to lead generation^15^. Within morphology analysis, cell painting is a popular method that entails analyzing fixed cells that are dyed with 6 stains to obtain information from 8 organelles and cellular components^16^, providing a generalizable, cost-effective framework for assessing perturbation affects. In parallel, image analysis has benefited from the development of machine learning techniques. One powerful, emerging machine learning technique is foundation modeling^17^, whereby a machine learning model is trained on a highly diverse data set enabling its application across a wide variety of tasks. The Deepcell system combines high resolution imaging of single cells in microfluidic flow with image analysis and feature extraction through the use of a cell morphology foundation model. This combination enables a label-free, non-labor-intensive means of analyzing and exploring diverse data sets, such as dissociated tumor cells (DTCs) and resistant cell lines across various cancer types and drug types.

In this study, we demonstrate with two case studies of drug-resistant cell populations, the utility of AI-driven single cell morphology analysis of high-resolution images to identify and classify resistant cells. This application has enabled us to identify morphological features that define the resistant cells across three different cancer types (ovarian cancer, breast cancer, and melanoma) and two different therapy types (chemotherapy, targeted therapy), further expanding the utility of image-based analyses. In three of the 6 resistant cell lines analyzed with the Deepcell system (A2780 Cisplatin Resistant, A2780 Doxorubicin Resistant, H460 Paclitaxel Resistant), there were pronounced changes in cell texture between the resistant and corresponding drug-naïve cells. In particular, the two A2780 resistant cell lines (Cisplatin, Doxorubicin) were characterized by a general loss of black blobs on the cell surface. This observation may be tied to a reduction of lysosomal compartments, which is characteristic of cisplatin-resistant A2780 cells and contributes to resistance through cisplatin sequestration away from nuclear DNA^18^. It may also point to similarities in the molecular mechanisms contributing to drug resistance to these two chemotherapeutics. Two of the melanoma cell lines (WM983B, 130425) exhibited pronounced increases in cell size between the resistant and naïve cells, where WM983B also exhibited a pigmented subpopulation; this increase in pigmentation is indicative of a melanocytic phenotype^19,20^ and has been shown to have protective features in the context of drug treatment^21^.

We envision that with expansion of the cell line panel for drug-resistance analysis, it will enable the extraction of morphology signatures that characterize specific resistance phenotypes, similar to the approaches applied to gene expression analysis. Analogously, we used heatmaps to effectively show morphological patterns in the data between sets of resistant and naïve cell populations, akin to a recent study in which morphology heatmap data of wild type and mutant cells from single gene CRISPR perturbations highlighted similarities between various ubiquitin-proteasome system mutants^22^. Another future aim will be to link particular morphology features and signatures to corresponding molecular pathways.

To emphasize the clinical potential of this study, we applied the H460-Paclitaxel resistance classifier to a dissociated tumor cell sample from a lung cancer patient and compared the resulting predictions to scRNASeq data from the same sample based on a gene signature of paclitaxel resistance in lung cancer^13^. We noted strong concordance between the percentage of cells predicted to be paclitaxel resistant by the classifier and the percentage of paclitaxel resistant cells based on gene expression. While this result is encouraging, future work will entail validating the cells predicted to be paclitaxel resistant by sorting these cells with the Deepcell instrument and using transcriptional and dose-response assays to confirm the phenotypes of the sorted cells. More generally, we envision that with further development of clinical-based classifiers, morphology analysis can be applied in a clinical context on tumor biopsies to enable personalized therapies; in other words, classifiers can be used on biopsies to understand which therapies might be most effective for a particular patient.

In sum, we show that morphology analysis through high resolution imaging and the development of foundation model-based classifiers is an effective approach for evaluating drug resistance in a label-free and non-labor-intensive manner. The approach has the potential to enable personalized therapies and complement extant methodologies (e.g. single cell transcriptomics) and drug development pipelines.

## Author contributions

Conceptualization: CR, SCB, MR, EL, MPL, MB, and AJ; Formal analysis: KS, ZL, RC, SS, MGM, AD, AJ; Methodology: CC, TP, VP, MGM, AD; Writing-original draft: CR, MB, AJ; Data curation: KS, ZL; Software: KS, ZL, SS; Writing-review and editing: CR, EL, MGM, AD, MPL, MB, AJ; Supervision: CR, MPL, MB, AJ; Resources: CR, EL, MPL, MB, AJ

## Declaration of interests

EL, MGM, and AD have no conflicts of interest. AJ, KS, ZL, CC, TP, RC, SCB, MR are current or former employees at Deepcell, Inc. SS, VP, MB, CR are current or former employees of Abbvie. MPL has unrelated research funding from Roche, Novartis, Molecular Partners, Oncobit, and Scailyte.

## Acknowledgments

The authors thank Mahyar Salek, Maddison (Mahdokht) Masaeli, Nianzhen Li, Kevin Jacobs, and Senzeyu Zhang for their contributions. BioRender was used for Fig. 1. This study received no funding.

