## Supplemental Information for "Deep learning-driven morphology analysis enables label-free classification of therapeutic agent-naive versus resistant cancer cells"

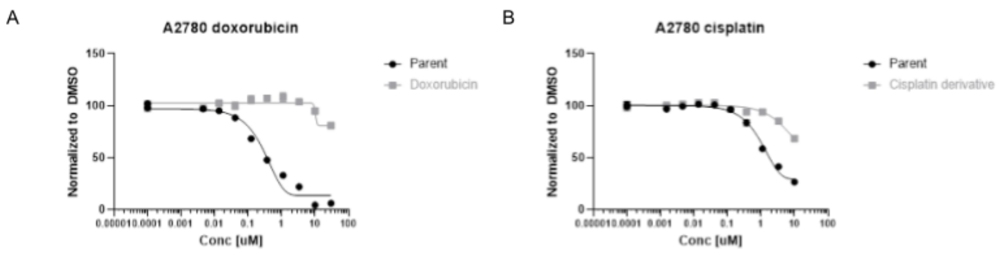


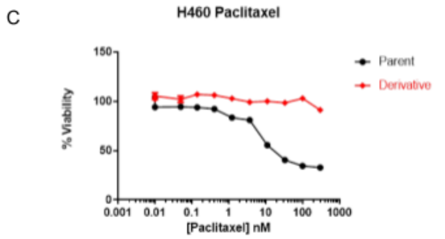


D

| Cell line | Drug | IC-50 (nM) |
|---|---|---|
| A2780 naive | doxorubicin | 0.35 nM |
| | cisplatin | 1000 nM |
| A2780 doxorubicin resistant | doxorubicin | >30,000 µM |
| A2780 naive resistant | cisplatin | >10,000 µM |
| H460 parent | paclitaxel | 29 nM |
| H460 paclitaxel resistant | paclitaxel | >300 nM |

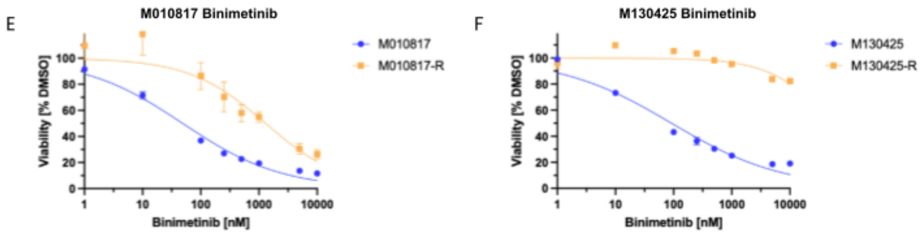


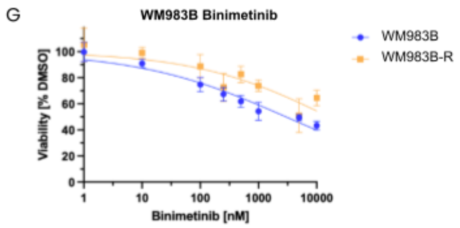


H

| Cell line | IC-50 (nM) |
|---|---|
| M010817 naive | 49 |
| M010817 resistant | 1303 |
| M130425 naive | 92 |
| M130425 resistant | 53139 |
| WM983B naive | 2972 |
| M130425 resistant | 16289 |

**Suppl. Fig. 1**: IC50 data for the A2780 and H460 resistant and naive populations used for training and validating the classifiers. A) IC50 curves for the A2780 doxorubicin resistant population (gray) compared to the naive population (black). B) IC50 curves for the A2780 cisplatin resistant population (gray) compared to the naive population (black). C) IC50 curves for the H460 cells paclitaxel resistant population (red) compared to the naive population (black). D) Table of the IC50 drug concentrations for each cell line/drug combination. E) IC50 curves for the M010817 binimetinib resistant population (orange) compared to the naive population (blue). F) IC50 curves for the M130425 binimetinib resistant (orange) compared to the naive population (blue). G) IC50 curves for the WM983B binimetinib resistant (orange) compared to the naive population (blue). H) Table of the IC50 drug concentrations for each cell line.

A

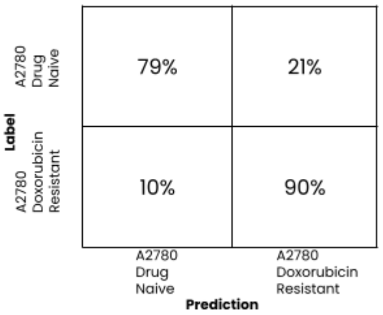


B

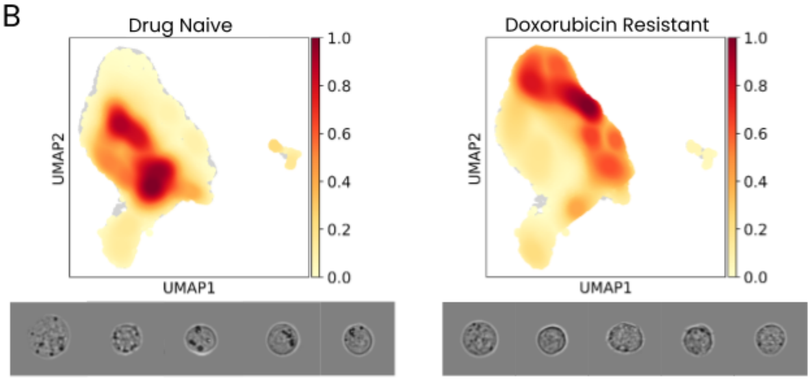


C

| Morphology Feature | Divergence Score |
|---|---|
| DL Feature 18 | 0.52 |
| DL Feature 62 | 0.52 |
| DL Feature 39 | 0.46 |
| DL Feature 10 | 0.45 |
| Large Black Blob, Integral | 0.43 |

D

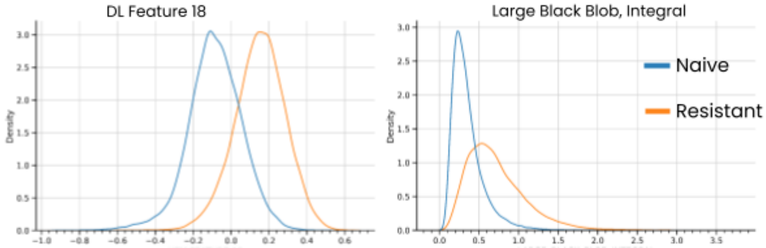


E

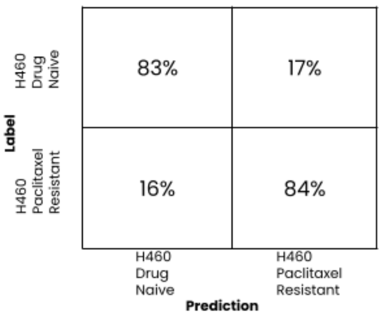


F

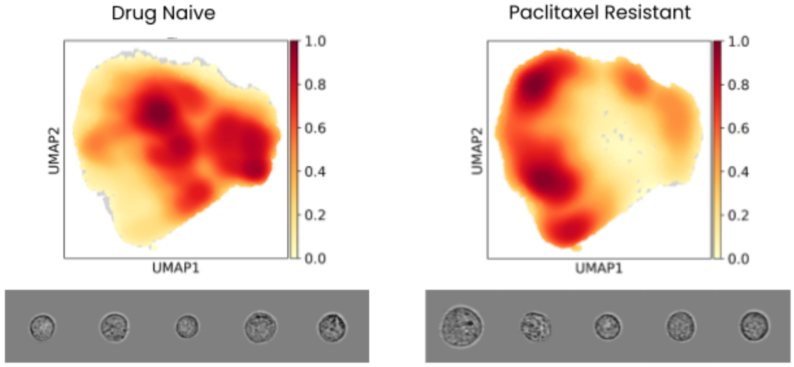


G

| Morphology Feature | Divergence Score |
|---|---|
| DL Feature 40 | 0.37 |
| DL Feature 55 | 0.36 |
| DL Feature 45 | 0.35 |
| DL Feature 24 | 0.34 |
| LBP Periphery 06 | 0.26 |

H

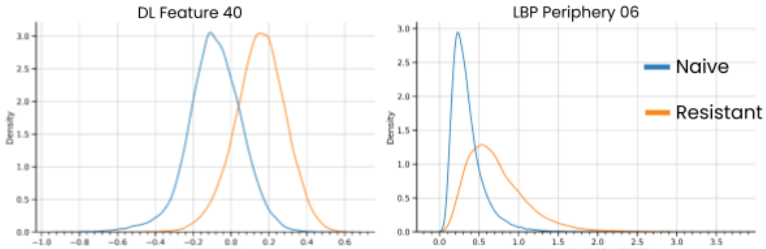

I

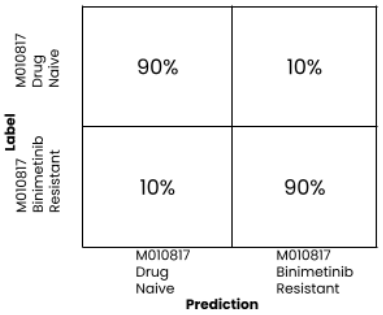


J

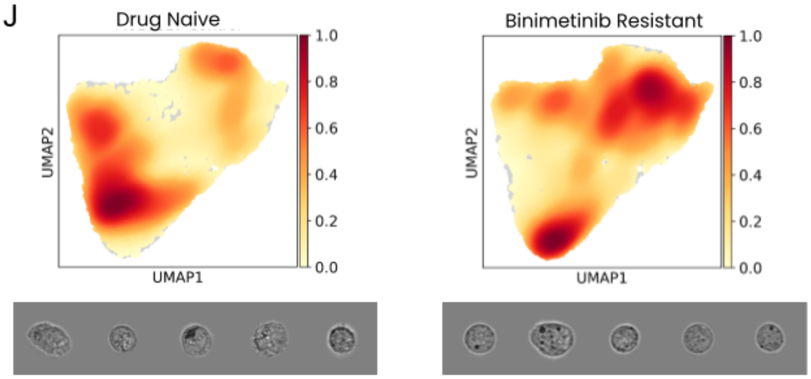


K

| Morphology Feature | Divergence Score |
|---|---|
| Percentile 25 | 0.41 |
| Negative Fraction | 0.37 |
| Mean | 0.35 |
| Hu Moment 1 | 0.35 |
| DL Feature 42 | 0.30 |

L

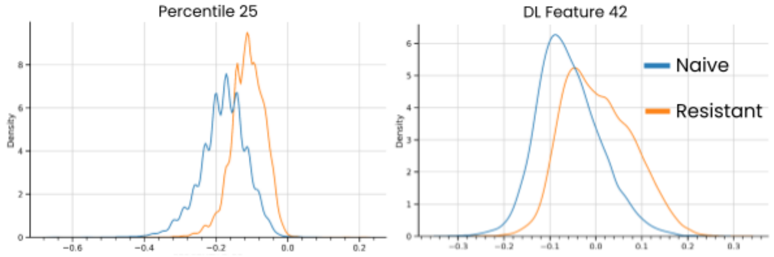


M

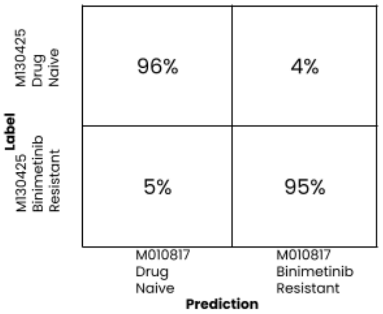


N

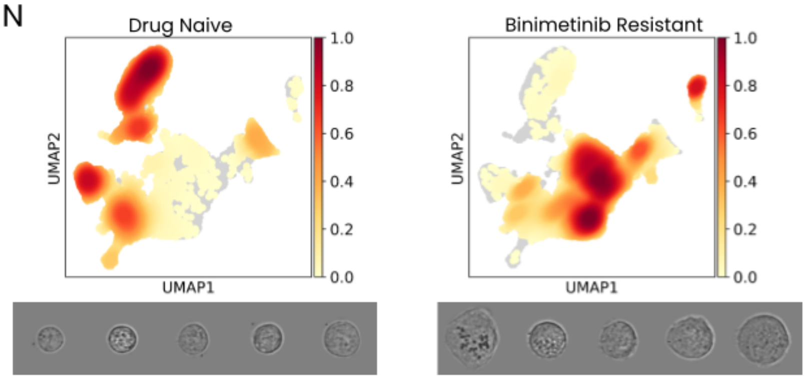


O

| Morphology Feature | Divergence Score |
|---|---|
| Max Feret | 0.60 |
| DL Feature 16 | 0.6 |
| Max Radius | 0.6 |
| Long Axis | 0.6 |
| Area | 0.59 |

P

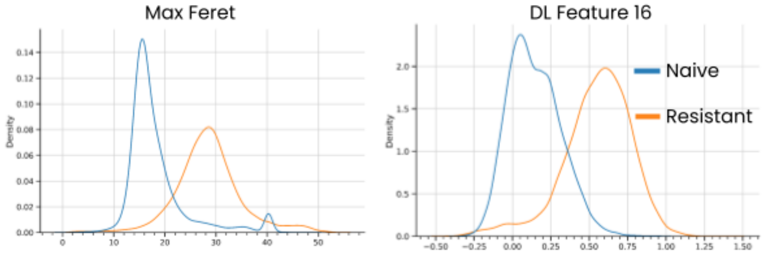

Q

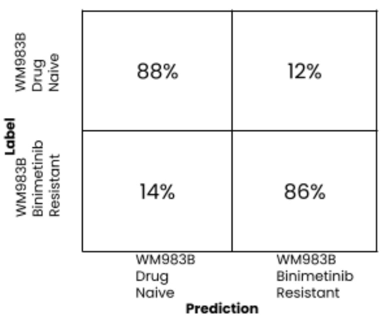


R

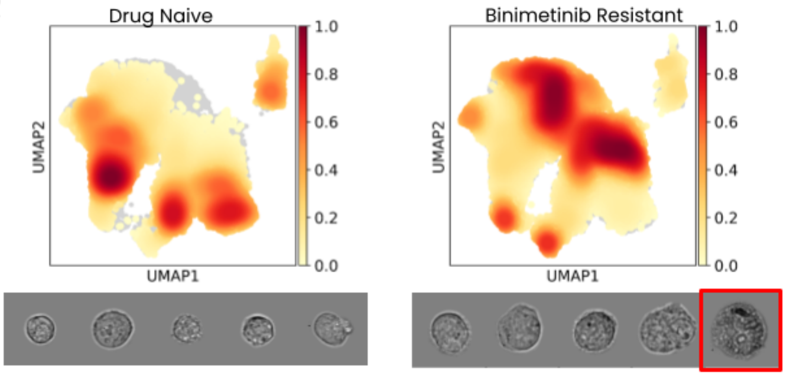


S

| Morphology Feature | Divergence Score |
|---|---|
| DL Feature 16 | 0.41 |
| DL Feature 13 | 0.39 |
| DL Feature 63 | 0.35 |
| DL Feature 19 | 0.35 |
| Min Radius | 0.31 |

T

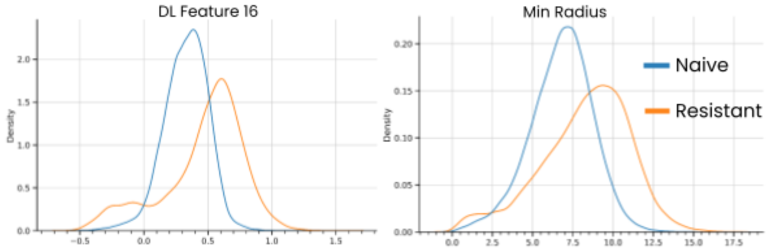


**Suppl. Fig. 2**: Morphology analysis of A2780 doxorubicin resistant and naive cells reveals key features differentiating the two populations enabling high accuracy classification based on morphology alone. A) Confusion matrix of the classifier for predicting Doxorubicin-resistant and naive A2780 cells based on morphology. B) UMAPs of the morphology features extracted by the Cell Morphology Foundation Model in density plot form, highlighting the morphological differences between the two populations. Representative images of each population beneath the respective UMAPs. C) Top 4 morphology features distinguishing the Cisplatin and Naive populations in A2780 cells, and the top morphometric feature in teal. D) Distributions of the top Deep Learning and Morphometric features between the naive and resistant populations. E) Confusion matrix of the classifier for predicting Paclitaxel-resistant and naive H460 cells based on morphology. F) UMAPs of the morphology features extracted by the Cell Morphology Foundation Model in density plot form, highlighting the morphological differences between the two populations. Representative images of each population beneath the respective UMAPs. G) Top 4 morphology features distinguishing the Cisplatin and Naive populations in H460 cells, and the top morphometric feature in teal. H) Distributions of the top Deep Learning and Morphometric features between the naive and resistant populations. I) Confusion matrix of the classifier for predicting Doxorubicin-resistant and naive M010817 cells based on morphology. J) UMAPs of the morphology features extracted by the Cell Morphology Foundation Model in density plot form, highlighting the morphological differences between the two populations. Representative images of each population beneath the respective UMAPs. K) Top 4 morphology features distinguishing the Cisplatin and Naive populations in M010817 cells, and the top deep learning feature in teal. L) Distributions of the top Deep Learning and Morphometric features between the naive and resistant populations. M) Confusion matrix of the classifier for predicting Doxorubicin-resistant and naive M130425 cells based on morphology. N) UMAPs of the morphology features extracted by the Cell Morphology Foundation Model in density plot form, highlighting the morphological differences between the two populations. Representative images of each population beneath the respective UMAPs. O) Top 5 morphology features distinguishing the Cisplatin and Naive populations in M130425 cells. P) Distributions of the top Deep Learning and Morphometric features between the naive and resistant populations. Q) Confusion matrix of the classifier for predicting Doxorubicin-resistant and naive WM983B cells based on morphology. R) UMAPs of the morphology features extracted by the Cell Morphology Foundation Model in density plot form, highlighting the morphological differences between the two populations. Representative images of each population beneath the respective UMAPs; an example of a pigmented cell is represented in the red box. S) Top 4 morphology features distinguishing the Cisplatin and Naive populations in WM983B cells, and the top morphometric feature in teal. T) Distributions of the top Deep Learning and Morphometric features between the naive and resistant populations.

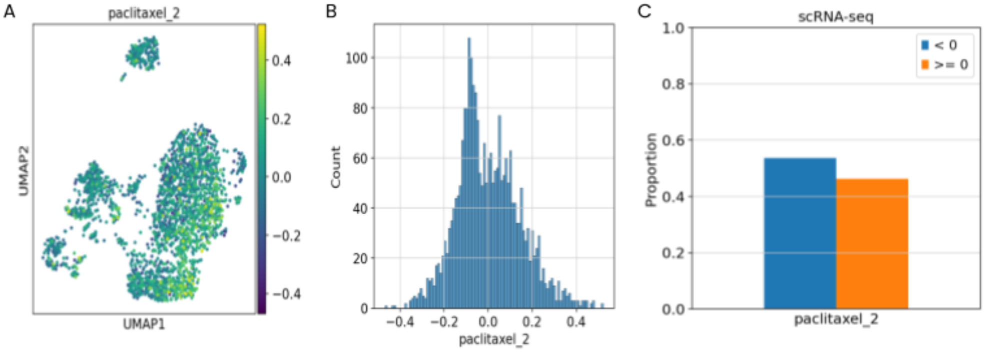


**Suppl. Fig. 3**: Transcriptional analysis of scRNASeq data phenotype analysis of the lung cancer DTC sample for comparison with the classifier predictions. A) UMAP of the scRNASeq data for the DTC sample constrained to the paclitaxel resistance gene signature. B) Histogram of cell counts for the expression levels of the paclitaxel resistance gene signature. C) Bar graph of the proportion of cells with normalized gene signature expression levels greater than (orange) and below 0 (blue). Cells with expression levels greater than or equal to 0 were considered resistant to paclitaxel.

| **Gene list** |
| --- |
| ATP6V0D1 |
| PKLR |
| SCLY |
| SMPD1 |
| DLC1 |
| PDPK1 |
| ZAK |
| STK24 |
| XPO1 |
| LMF1 |
| FAM164A |
| CCDC99 |

**Suppl. Table 1**: List of genes comprising the paclitaxel resistance gene signature, taken from Ref. [13].
